# A temperature-controlled, circular maintenance system for studying growth and development of pelagic tunicates (Salps)

**DOI:** 10.1101/2023.07.21.547827

**Authors:** Svenja J. Müller, Wiebke Wessels, Sara Driscoll, Evgeny A. Pakhomov, Lutz Auerswald, Katharina Michael, Bettina Meyer

## Abstract

Salps are pelagic tunicates that are able to form large blooms under favorable conditions by alternating between sexual and asexual reproduction. While their role in the regional carbon cycle is receiving attention, our knowledge of their physiology is still limited. This knowledge gap is mainly due to their fragile gelatinous nature, which makes it difficult to capture intact specimens and maintain them in the laboratory. We present here a modified kreisel tank system, that was tested onboard using the Southern Ocean salp *Salpa thompsoni* and station-based using the Mediterranean species *Salpa fusiformis*. Successful maintenance over days to weeks allowed us to obtain comparable relative growth and developmental rates as *in situ*, and provided insight into their potential life cycle strategies. By providing a starting point for successful maintenance, we hope to stimulate future experimental research on this understudied taxonomic group.

## Introduction

In recent decades, gelatinous zooplankton received an increased attention as it frequently dominates global marine zooplankton communities worldwide, arguably as a result of overfishing and anthropogenic climate change (Richardson *et al*., 2009). Research mainly focused on true jellyfishes (cnidaria) and ctenophores (comb jellies) so far, whereas pelagic tunicates, salps in particular, were generally overlooked (Henschke *et al*., 2016).

Salps are found in open ocean waters with the exception of the Arctic (Madin and Deibel, 1998). They are known as efficient filter feeders and frequently occur in dense aggregations, when food is abundant (Madin *et al*., 1996; Andersen, 1998). The development of these so-called blooms is facilitated by the high reproductive capacity of salps during the asexual stage (i.e., oozoids, solitaries) alternating with the sexually reproducing blastozooids (i.e., aggregates) (Foxton, 1966; Madin *et al*., 1996; Madin and Deibel, 1998). Salps are able to displace other zooplankton on small spatial scales, can play significant roles in regional biogeochemical cycling (Fischer *et al*., 1988; Lee *et al*., 2010; Gleiber *et al*., 2012; Böckmann *et al*., 2021; Pauli *et al*., 2021) and serve as food sources for higher trophic levels (Harbison, 1998; Henschke *et al*., 2016).

Over the past decades, global warming led to dramatic changes in abundance and distribution of salps, which is supposed to have profound implications for the ecosystem (Loeb *et al*., 1997; Atkinson *et al*., 2017). Currently, the role of salps in marine ecosystems is increasing, but only little is known about their ecology, physiology and life cycle (Henschke *et al*., 2016). This restricts our understanding of marine carbon export and therefore, its adequate implementation in regional and global biogeochemical models. In addition, the lack of knowledge of salp biology and their sensitivity to future climate change scenarios makes it difficult to predict future population dynamics and to assess the potential impact of a salp-dominated zooplankton community on an ecosystem. This knowledge gap can be mainly explained by the seasonal and spatially patchy occurrence of salps that only allows for narrow time windows for research, where especially the latter making them a challenging research subject. Furthermore, their fragile gelatinous nature restricts field sampling of intact specimens with nets or trawls and makes them delicate organisms for long-term maintenance in aquaria for experimental studies (Henschke *et al*., 2016).

In first attempts, individuals of the pelagic tunicate *Thalia democratica* were kept in the laboratory in jars filled with daily renewed seawater (Braconnot, 1963). This method resulted in high mortalities because the salps became entangled to the surface film. This issue was resolved in a subsequent study by Heron (1972) using small tanks with lids. A detailed description of the cultivation technique is lacking, however, and it is doubtful that the growth rates of *T. democratica* measured by Heron (1972) were unbiased: The initial phytoplankton was grazed down during the experiment. A cultivation technique introduced by Paffenhöfer (1970) used a gentle stirrer and was successfully tested on the appendicularians *Oikopleura dioica* and *Fritillaria borealis*, two small (∼1 mm) representatives of pelagic tunicates (Paffenhöfer, 1973, 1976; Paffenhöfer and Harris, 1979). However, the most useful aquarium for maintaining pelagic gelatinous zooplankton seems to be the planktonkreisel, first introduced as a horizontal device by Greve (1968) and modified by Greve (1975) and Hamner (1990). The cultivation of gelatinous zooplankton in the planktonkreisel was successfully tested and described for true jellyfishes and ctenophores (Greve, 1970, 1975; Raskoff *et al*., 2003; Courtney *et al*., 2020; Lechable *et al*., 2020; Ballesteros *et al*., 2022). For salps, the first maintenance attempts in kreisel tanks have only been published in recent years (Lüskow *et al*., 2020; Stukel *et al*., 2021). However, these studies only involved short-term incubations of salps (∼ 24 h, Stukel *et al*., 2021) and lacked more detailed maintenance descriptions (Lüskow *et al*., 2020; Stukel *et al*., 2021). This limits implementation of physiological (long-term) studies to qualify and quantify salp life strategies, and to obtain reproducible physiological rates for parameters such as e.g., growth, oxygen consumption, and development under controlled experimental conditions. In addition, studies on their response to changing environmental conditions are also limited, which is particularly important in the light of ongoing climate change.

The objective of our study was to construct a temperature-controlled circular and flow-through aquarium system for salps in order to facilitate future studies on salp physiology and their response to changing environmental conditions. We tested our system by maintaining two salp species from two different habitats: *Salpa thompsoni* from the Southern Ocean, and *Salpa fusiformis* from the Mediterranean. The experiments with *S. thompsoni* were conducted during a ship-based expedition onboard the German research vessel *Polarstern* in the Southern Ocean, whereas the experiments with *S. fusiformis* took place at the Oceanological Observatory in Villefranche-sur-Mer (France). In addition to a detailed description of our aquarium-system, we highlight limitations, improvements and advice for field sampling and future experimental studies on salps. We present results on growth and developmental rates of both salp species maintained in our experimental kreisel tank systems, validated by *in situ* rates and published data.

## Materials and Procedures

### Kreisel tank sets and custom modifications

The kreisel tank is a well-established aquarium for the maintenance of gelatinous zooplankton (Greve, 1970, 1975; Raskoff *et al*., 2003; Courtney *et al*., 2020; Lechable *et al*., 2020; Ballesteros *et al*., 2022). To maintain the different salp species (*S*. *thompsoni* and *S. fusiformis*) in our study, two sizes of the water-running breeding kreisel tank (Schuran Seawater Equipment, Jülich Germany) were used:

I. V= ∼73.10 l, diameter= 610 mm, depth= 250 mm (here after big kreisel tanks, *n=*6)
II. V= ∼14.14 l, diameter= 300 mm, depth= 200 mm (here after small kreisel tanks, *n=*6)

The big kreisel tanks (I) were connected in pairs, whereas the small kreisel (II) tanks in a set of three, each of the two set ups sharing one outflow (Figure 1A, B). However, the inflow of the individual kreisel tanks could be controlled independently by valves attached to each inlet pipe. The larger salp species, *S. thompsoni*, was maintained only in the bigger kreisel tanks (I), while for the smaller Mediterranean species *S. fusiformis* both kreisel sets (I, II) were used (Table 1).

**Figure 1.**
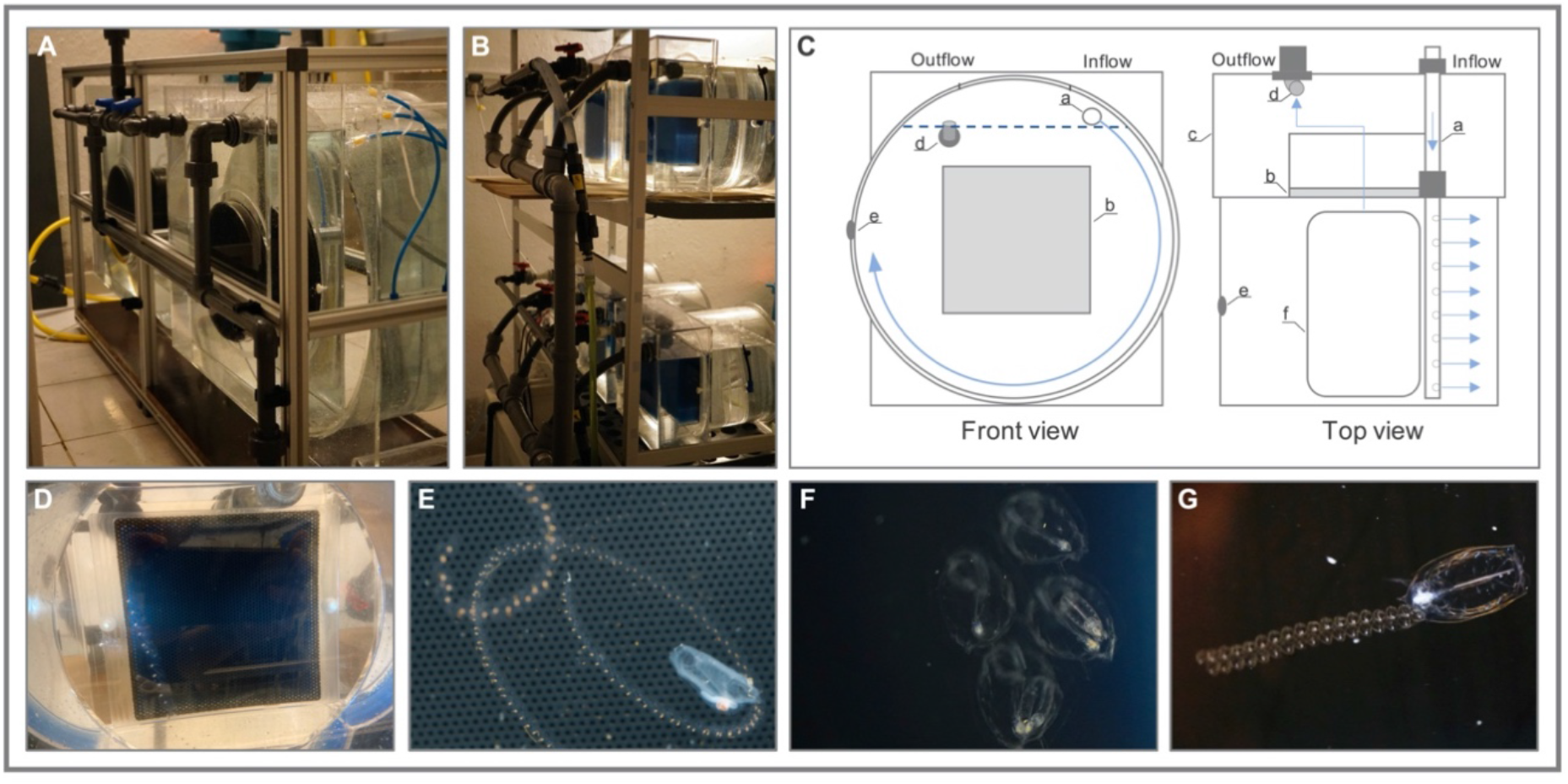
Experimental kreisel tank set up and salps maintained. (A-B) Set up with big kreisel tanks, each connected in pairs ((A), V= 73.10 L) and small kreisel tanks each connected in a set of three ((B), V=14.14 L). (C) Schematic representation of the functionality of the kreisel tank used in front and top view: a=spray bar, b=filter (pore size: 2mm), c=external filter chamber, d=elbow tubes in outflow pipe, e=air connection valves, f=lid. The light blue arrows indicate the direction of water flow. The dashed horizontal line marks the water level. (D-E) Images of chain of aggregates (D, E) and a single solitary (E) of *S. fusiformis* maintained in the kreisel tank system. Image of a chain of aggregates (F) and a solitary in the progress of releasing a chain of aggregates (G) of *T. democratica* in our kreisel tank system. Photo credit: (A-B), (D), (F-G) S.J. Müller, (E) A. Jan; 2021, Oceanological Observatory of Villefranche-sur-Mer, France.

**Table 1.**
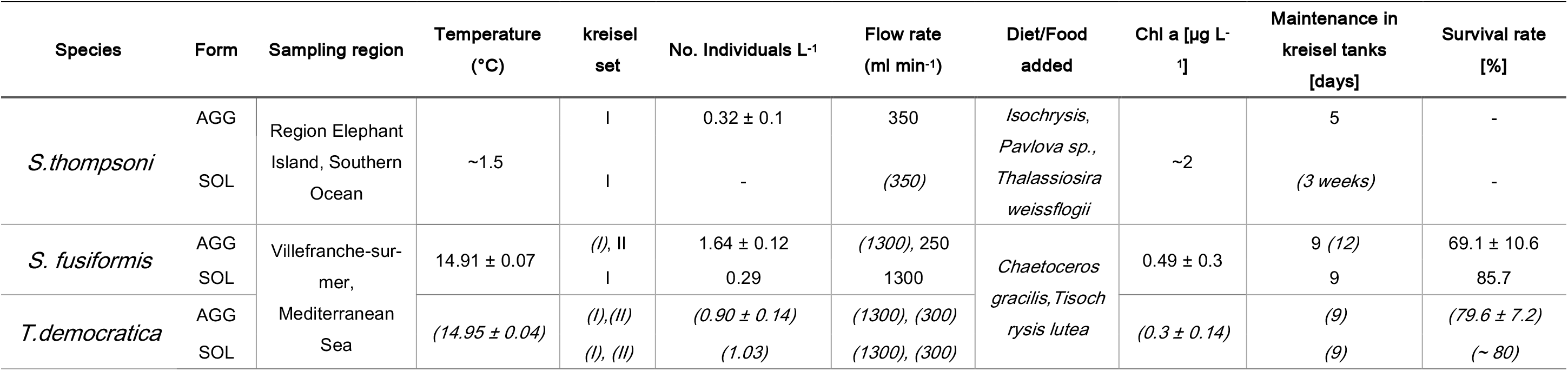
Overview of the maintenance of *S. thompsoni, S. fusiformis* and *T. democratica* in kreisel tanks. Information’s are presented species and form-specific (AGG= aggregate, SOL= solitary). Breeding kreisel set refers to the I) big kreisel tanks (V= ∼73 l) and II) small kreisel tanks (V= ∼14 l). All data refer to maintenance settings during the experiments determining growth and development rates. Data in brackets and italics refer to settings for successful cultivation attempts outside the experiment.

Although the kreisel tanks differed in size, the general technical functionality was similar (Figure 1C): The inlet pipe (spray bar, Figure 1C, a) in each kreisel tank created a uniform circular flow that prevented organisms from any contact with the water surface or tank walls (Greve, 1968; Raskoff *et al*., 2003). The water flowed in a circular stream and drained through a centrally positioned filter (pore size: 2 mm, Figure 1C, b) located in the back of the tank into the external filter chamber (Figure 1C, c). A sponge located behind the filter reduced the water suction flow from the kreisel into the filter chamber. Moreover, the following custom modifications were made: the filter holes were deburred to prevent the tunic of salps from tangling on the sharp edges of the holes. For the small kreisel tanks, the water level was raised to the level of the spray bar by installing elbow tubes in the outflow pipe (Figure 1C, d). All kreisel tanks were additionally equipped with air connection valves (Figure 1C, e) to optionally switch to an air-running system. As an additional modification for the usage of the big kreisel set on ship-based expeditions, an aluminum frame and a floor mount were built to provide protection against ship motion (Figure 1A). The position of each spray bar was adjusted wall-sided to prevent bubble formation. A full list of the equipment used can be found in Supplementary Table S1. Further details on the specific experimental set up for each salp species is given below.

### Field sampling and laboratory maintenance of S. thompsoni

#### Sampling

For onboard experiments, *S. thompsoni* were caught during expedition PS112 with R.V. *Polarstern*, close to Elephant Island (60°54.777’S 054°51.036’W) on April 24^th^, 2018, using an Isaacs-Kidd mid-water trawl (IKMT 1.8 m^2^ mouth opening, 550 μm mesh size) (Figure S1A). To obtain individuals in healthy condition, still pumping and swimming, the upper 50 m were sampled at a maximum speed of up to 2.0 knots, employing a closed cod end, equipped to the net. Immediately upon capture, salps were sorted by forms (aggregates and solitaries), and chains of aggregates. Chains (*n=*19) of different sizes and length (3-9 individuals chain^-1^) were carefully transferred into big kreisel tanks (I, *n=*5) shortly after they were caught. Sea surface temperature (SST) during sampling was ∼ 0.51°C.

In vicinity of Deception Island (⁓ 62°34’S – 59°55’W), *S. thompsoni* were collected at the multi-day station, March 27 to April 1, 2018 using oblique IKMT tows in the top 0-170 m water layer in order to obtain field sample abundance and growth (see Wessels *et al*., 2018).

#### Maintenance

Animals were kept in a temperature-controlled room for 5 days under natural/local light conditions (13h:11h/day:night) and fed three times a day with a mix of different instant algae (*Isochrysis*, *Pavlova sp., Thalassiosira weissflogii)* at a chlorophyll concentration of ∼2 µg L^-1^ (Iso 1800^TM^, PAV 1800^TM^, TW 1800 ^TM^; Reed Mariculture, CA, USA). Filtered (5 μm) ambient seawater was pumped through the kreisel tanks at a constant flow rate of ∼350 ml min^-1^ using Eheim pumps (Eheim, Universal 1200, Germany). Due to the passage through the vessels’ pump system, the temperature of the filtered seawater was ∼ 1.5 °C when arriving at the kreisel tank set up. A turnover rate (defined as the time needed to fully replace the water per tank) of ∼ 6.5 h ensured sufficient distribution of food and prevented accumulation of harmful bacteria. Coarse impurities were removed daily with a glass tube. Salinity and temperature were checked regularly using a Conductometer WTW 3110 (WTW, Germany).

### Field sampling and laboratory maintenance of S. fusiformis

#### Sampling

Field sampling was conducted by snorkeling during a salp bloom period in spring (March/April 2021) at sunrise, using ziploc-bags or a beaker when salps were near the surface (Figure S2). The salps in the sampling devices were stored in a cooler box to keep water temperature constant during transport. After checking for parasites (amphipods, copepods), healthy specimens were transported to the kreisel set-up in the temperature-controlled room (16°C) at the Oceanological Observatory, Villefranche-sur-mer, France, within 1.5 hours after capture.

During the same bloom period, salps were sampled onboard R.V. *Sagitta* at regular intervals at Point B at the mouth of the bay of Villefranche-sur-Mer (Figure S1B). This was to estimate the abundance and growth rates of field samples using a Regent net (1 m^2^ mouth opening, 680 μm mesh size, towed obliquely) in the upper 80 m. The filtered volume was estimated by attaching a flow meter placed close to the center of the net. Upon retrieval of the nets, subsamples were preserved in 4% formaldehyde for further analyses at the Alfred-Wegener-Institute Helmholtz Centre for Polar and Marine Research, Bremerhaven. The *in-situ* conditions at Point B were characterized by a mean Chl a concentration of 0.24 ± 0.1 μg L^-1^ and temperature of 14.82 ± 0.30 °C throughout the whole sampling period (Service d’Observation en Milieu Littoral; www.somlit.fr).

#### Maintenance

For the maintenance of aggregates and solitaries of *S. fusiformis* in the laboratory we used a kreisel-tank system that consisted of one set of big kreisel tanks (I, total *n=*2) and two sets of small kreisel tanks (II, total *n=*6), and a reservoir (V= ∼ 200L). While solitaries of *S. fusiformis* were only maintained in big (I) kreisel tanks, aggregates of *S. fusiformis* were maintained in big (I) and small (II) kreisel tanks (Figure 1 D, E, Table 1). Animals were kept under a natural day: night rhythm of 11h:13h and were fed with an algal mix of 50 ml cultured *Chaetoceros gracilis* and 50 ml *Tisochrysis lutea*, which was added to the reservoir daily. This experimental set up resulted in a constant water temperature of ∼ 15 °C and a Chl *a* concentration of 0.49 ± 0.3 μg L^-1^, similar to the *in-situ* conditions at sampling Point B (outlined above). To ensure a stable water temperature of 15 °C, an inline aquarium cooler (TK3000, TECO) was employed that circulated the water in the reservoir by a submersible pump (Eheim, Universal 2400, Germany). Filtered seawater (5 μm cartridge filter) was continuously supplied into the ∼ 200 L reservoir, with a flow rate of 4200-4800 ml min^-1^. Flow from the reservoir to each big and small kreisel tank was maintained with pumps (Eheim, Universal 1200, Germany) at a flow rate of 1300 ml min^-1^ and 250 ml min^-1^, respectively. Water from the small kreisel tanks flowed back into the reservoir, while water from the large tanks was discharged directly and was replaced by new incoming filtered seawater. In this way, a turn-over rate of approx. 3 hours (V= approx. 430 l) was achieved at constant temperature. Water temperature was monitored by a logger (HOBO pedant temp/light logger (MX2202), Onset, MA, USA), placed into the overflow chamber of a randomly selected kreisel tank system. Water level was controlled by an overflow pipe connected to the reservoir. After recording the whole animal performance and sampling, coarse impurities and dead individuals were carefully removed daily using a glass tube.

### Determination of relative growth and developmental rates

#### S. thompsoni

Immediately after field sampling, one individual salp from each chain (*n=*19) was carefully separated by a spring-steel tweezer. Oral-atrial length (OAL) was measured and each specimen was staged according to Foxton (1966) before transferring the rest of each chain to the kreisel tanks (*n=*5). All individuals in a chain remained connected and were maintained under constant feeding conditions at ∼1.5 °C for 5 days. On the fifth day after capture, all individuals (individuals chain^-1^ *n=*3-8) of the respective chains were sampled.

For determination of *in situ* growth of aggregates, a subsample of > 200 salps was taken during the multi day-station. All individuals were measured for OAL with an accuracy of 1 mm, for use in cohort analysis (Wessels *et al*., 2018).

#### S. fusiformis

After field sampling, 6 chains of aggregates were separated into halves by gently twisting a spring-steel tweezer back and forth, before transferring one half into the kreisel tank system. The resulting 6 chain halves were distributed in the 6 small kreisel tanks (1 chain kreisel^-1^). There were no solitaries of *S. fusiformis* found during any of the morning samplings in the bay of Villefranche-sur-mer, France. As an alternative of tracking growth of solitaries, a more developed chain of aggregates, almost all carrying a fully developed solitary, was placed in a big kreisel tank. After three, seven and nine days spent in kreisel tanks, individuals (*n=*6-7) of *S. fusiformis* of both forms were sampled from each kreisel tank. Based on the similarity of embryonal and stolon development to *S. thompsoni, S. fusiformis* was staged according to Foxton (1966) under a microscope (Nikon C-LEDS, No. 224659, China) equipped with a digital camera (Dino-Eye AM7025X, USA) followed by measurements of OAL and total length (TL) at each sampling.

*In situ* growth of aggregates of *S. fusiformis* from the field was measured from subsamples of paraformaldehyde field samples (see above), removed by a Henson-Stempel pipet (50 ml). Size of the subsample was chosen depending on the density of salps. Samples were counted, and OAL and TL of both forms of *S. fusiformis* were measured and staged according to Foxton (1966). A 10% shrinkage correction was applied due to effects of formalin (Heron *et al*., 1988). Estimation of growth rates using cohort analysis were calculated under the assumption of sampling the same salp population.

### Statistical analysis and data visualization

Relative growth rates (% body length day^-1^) in the kreisel tanks were calculated by combining the OALs of all individuals measured during first sampling (*OAL*_O_) with OALs of all individuals measured at the end of the experiment (*OAL_t_*) within each chain (aggregates) or within siblings (solitaries). Each individual relative growth rate was then determined as:

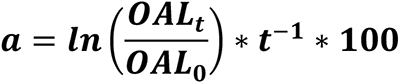

This cross-combination of individuals within chains/siblings was done to account for the uncertainty in initial OAL of the salps sampled at the end of the experiment. The approach allowed us to obtain growth rate ranges instead of point estimates, providing a more robust metric for comparison of the effects of initial stage and initial length on the rate. A linear regression model was used to explore the effect of mean initial OAL on relative growth rate for each chain/group of solitaries.

R package mixtools (Benaglia *et al*., 2009) was used to detect cohorts from the field samples, mean OAL ± SD were extracted and relative growth rates were estimated according to the equation stated above.

To estimate developmental rates in the kreisel tanks (i.e., the time [in days] required to develop to the next stage) the mean stage of all individuals within a chain/ all solitaries was calculated during the first sampling (*Stage_O_*) and at the end of the experiment (*Stage_t_*).

The mean development rate for each chain/solitaries was defined as:

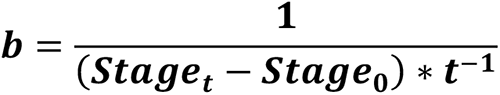

Aggregates of stage “X” (i.e., stage not clearly identified) were neglected in this calculation. Mean and standard deviation (SD) of development rates were obtained by grouping variable initial stage.

The environmental data (temperature, Chl a) in Villefranche-sur-Mer (Point B) were extracted from the SOMLIT database (Service d’Observation en Milieu Littoral; www.somlit.fr) on [30.06.22]. All data were processed and visualized with help of *tidyverse v*. 1.3.2 (Wickham et al., 2019).

### Assessment

#### Suitability of the kreisel tank system for maintenance of salps

Kreisel tanks have been used for breeding attempts of true jellyfishes and ctenophores (Greve, 1970, 1975; Raskoff *et al*., 2003; Courtney *et al*., 2020; Lechable *et al*., 2020; Ballesteros *et al*., 2022). However, the suitability of the circular kreisel tank for maintaining salps has never been described. In our study, the circulation of the kreisel tank gently directed salps away from the tank surface by a circular flow, preventing collision. We also observed no evidence of damage due to wall contact. By adjusting the water level in the kreisel tanks (see *Kreisel tank sets and modifications*), we were able to reduce harmful bubble formation in the tanks and did not observe any specimens getting caught at the water surface. The flow rate of each kreisel was adjusted to meet the different needs of any species and form without causing them to tumble (Table 1).

While some jellyfish species require different breeding tanks for different stages (Lechable *et al*., 2020; Ballesteros *et al*., 2022), our kreisel tanks appeared to be suitable for culturing almost all stages of salps. We were able to maintain *S. thompsoni* and *S. fusiformis* in several trials ranging from days to weeks by adjusting the system to the needs of the respective species and form (Table 1). The system was also successfully tested with the smaller Mediterranean species, *Thalia democratica,* at the Oceanological Observatory of Villefranche-sur-Mer in spring 2021. Both forms (aggregates and solitaries) were maintained up to 9 days in the kreisel tank system, underlining the versatility of our system (Figure 1F, G). Summarized information on maintenance of *Thalia democratica* is given in Table 1. However, due to a small number of individuals sampled and a low number of buds chain^-1^, *Thalia democratica* were excluded from the growth and developmental rate analysis here. Although the kreisel tanks appeared to be suitable for all salps and forms tested, two limitations should be noted at this point and considered in future experiments: With the exception of *S. thompsoni* and solitaries of *S. fusiformis*, individuals of the different species were maintained during culturing attempts in both kreisel tank sizes (I, II, Table 1). Due to the larger size of *S. thompsoni* and the strong swimming behavior of the solitaries of *S. fusiformis*, large kreisel tanks (I) are recommended for future maintenance experiments for both. Moreover, newly released chains of the smaller species, *T. democratica*, with a total body length of ∼2 mm tended to break and were occasionally sucked through the 2 mm pores of the filter. It is therefore advisable, to use a filter (Figure 1C, b) with 1 mm pores and reduce the flow rate, at least until the animals have grown beyond a minimum size of 3-4 mm. During maintenance, successful releases of chains by asexual reproduction and of young solitaries by sexual reproduction of both Mediterranean species were observed (personal observation, Figure 1G, Supplementary Video 1). This is proof of the favorable rearing conditions in our kreisel tanks.

By using a flow-through system, we ensured high water quality and a continuous natural food supply through filtered seawater (5 μm) in combination with instant algae and phytoplankton cultures for *S. thompson*i and both Mediterranean species (*S. fusiformis*, *T. democratica*), respectively. The described kreisel tanks and their custom modifications (see *Kreisel tank sets and modifications*) would also allow switching to a closed system, in combination with biological filtration and UV disinfection. In terms of experimental studies, our system is fully adjustable and allows for different experimental designs (e.g., temperature manipulation), that will be crucial for improving predictions on the performance of salps in the face of global climate change. Consequently, different temperature regimes were tested at the Oceanological Observatory of Villefranche-sur-Mer in spring 2021: By using a titanium heater (500 W, Aqua Medic, Germany) in combination with a temperature controller (Aqua Medic T-controller twin, Germany) placed in the reservoir, we were able to perform a stable temperature increase from 15 °C to 17 °C and 15 °C to 18 °C by a steady temperature increment of 1 °C day^-1^ and 1.5 °C day^-1^, respectively. Target temperature was kept stable at ± 0.2 °C for several days, indicating a reliable system for future experimental designs.

#### Growth and developmental rates of S. thompsoni and S. fusiformis

Most studies examining growth and development of salps are based on modeling approaches (Henschke *et al*., 2018; Groeneveld *et al*., 2020) and cohort analyses (Loeb and Santora, 2012; Pakhomov and Hunt, 2017; Lüskow *et al*., 2020). Rates published so far are therefore mostly based on short-term observations and vary between regions and/or studies. By providing a circular kreisel tank system as a potential future tool for monitoring salp performance, growth and development, we were able to gain novel insights into potential life cycle strategies of salps. In the following, growth and developmental rates of aggregates of *S. thompsoni* and of both forms (aggregates and solitaries) of *S. fusiformis* that were maintained in our kreisel tank system were calculated and evaluated using rates determined *in situ* at each study site. Furthermore, the results are related to the existing knowledge of growth and development rates of *S. thomspsoni* and *S. fusiformis*.

#### S. thompsoni

Chains (*n=*19) of aggregates of *S. thompsoni* were maintained in big (I) kreisel tanks (*n=*5) onboard R.V. *Polarstern* during PS112 for 5 days (Figure 1A). The focus of this experiment was on comparing growth and development within chains (representing clones). The duration of the experiment was therefore limited by the number of individuals per chain and number of samplings.

The size of aggregates appeared to vary within a chain (± SD range 0-1.7 mm, Figure S3A, B), so mean relative growth rates (% body length day^-1^) were determined by combining the OAL of all sampled individuals within a chain. The mean relative growth rates estimated from change in body length ranged from -1.64 to 3.15 % d^-1^(-0.36-0.36 mm d^-1^) (Figure 2A). In comparison, *in situ* growth rates assessed near Deception Island using cohort analysis were ⁓ 5% d^-1^ (⁓ 0.4 mm d^-1^) for 8 mm aggregates (Wessels *et al*., 2018). This growth rate corresponds to the upper end of the growth estimates in our experiments, probably due to the neglect of negative growth in cohort analyses. Upper growth rates in this study (0-3.15 % d^-1^) were within the rates calculated for *S. thompsoni* caught near the Antarctic Peninsula of 0.3-4.6% d^-1^ (Loeb and Santora, 2012), but in the lower range of *in situ* rates for aggregates from the Antarctic Polar Front zone (3.7-9.1% d^-1^) applying cohort analysis (Pakhomov and Hunt, 2017). In a study by Lüskow *et al*. (2020), a newly released (⁓ 5 mm OAL) chain of aggregates from the Chatham Rise region, New Zealand, was held in a kreisel tank for 5 days at 11 °C and high growth rates of 8.78 % d^-1^ were observed. These were comparable to *in situ* growth rates (9-11% d^-1^) of ⁓ 10 mm OAL aggregates obtained using cohort analyses in the same study (Lüskow *et al*., 2020). The differences in growth rates between the studies can be explained by differences in temperature, size and food concentration in the different regions, which both are known to affect salp growth rates (Heron, 1972; Deibel, 1982). Therefore, the lower SST of -1 to 2 °C in the study by Loeb and Santora (2012) and of ∼1.5 °C in our study may explain the lower growth rates compared to the higher growth rates observed for salps from the chlorophyll a rich APF region at ∼4-5 °C (Pakhomov and Hunt, 2017) and in the kreisel tank at ∼ 11 °C in the study by Lüskow *et al*. (2020).

**Figure 2.**
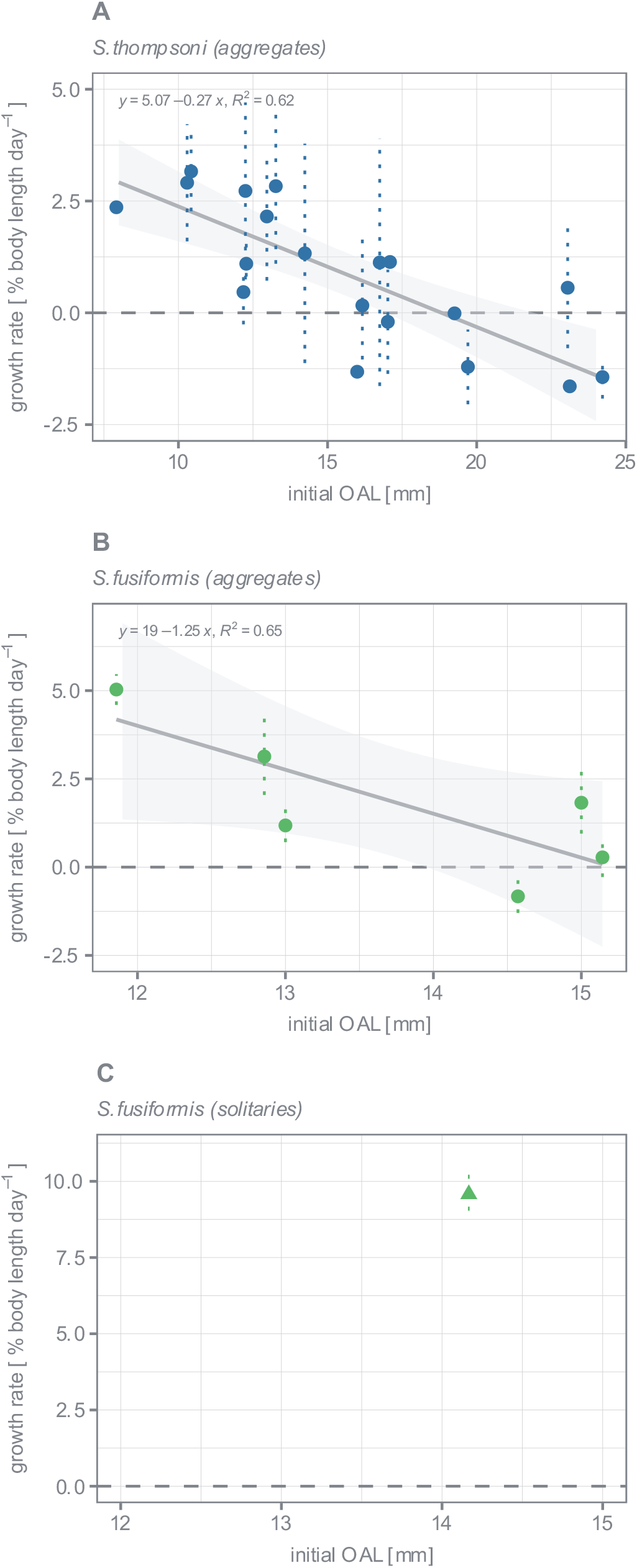
Relative growth (% body length per day) rates of *S. thompsoni* and *S. fusiformis*. Mean growth rates and confidence interval of individuals cross-combined within chain of aggregates (A, B) or group of solitaries (C) were calculated as a function of initial mean OAL for aggregates of *S. thompsoni* (A), and for aggregates (B) and solitaries (C) of *S. fusiformis*. Regression model and 95 % confidence interval are shown. The dashed horizontal line marks the putative threshold for the increase/decrease of daily rates. Each data point represents the mean growth rate of a single chain of aggregates (A, n=19; B, n=6) or a group of solitaries (n=8).

The negative growth rates observed in our study were only found in chains with a larger initial OAL (range of 16-24 mm). However, it is questionable whether the salps began to shrink after reaching a certain OAL (and or developmental stage) or merely reduced their growth rates for a short period of time. Nevertheless, the relative growth rate decreased moderately with increasing initial OAL, yielding a regression function of y= 5.07-0.27x (R^2^=0.62, Figure 2A), a trend that was already observed for aggregates of *S. thompsoni* using cohort analyses (Pakhomov and Hunt, 2017; Lüskow *et al*., 2020; Henschke *et al*., 2021). An increased growth rate at the beginning of life may be beneficial to outgrow the feeding range of most planktonic predators (Heron, 1972). A reduced energy investment into somatic growth might also be possible (and potentially shrinking in variable and unpredicted environments). This would favor embryonic development at a certain stage, as the embryonic growth depends on a continuous supply of nutrients from the parental aggregate via the placenta (Chiba *et al*., 1999). For other gelatinous organisms, such as ctenophores and medusae, a shrinkage was observed under poor conditions in favor of keeping essential metabolic rates unchanged (Harbison, 2009). The potential ability of aggregates to shrink at advanced stages could be a strategy to successfully develop their offspring at a maximum rate regardless of external circumstances. Our results suggest that this strategy of energy conservation may begin early after fertilization (stage 1.5, Figure S4) and becomes more evident as the embryo continues to develop within the parental aggregate. About 81.5 % of all individuals that showed potential negative growth rates were of initial developmental stage 2 or higher at the first sampling timepoint (Figure S4A). However, our data does not allow to distinguish between the effects of increasing OAL and initial stage on the relative growth rate. Although the developmental stage of salps, defined by the gradual growth of the embryo inside, depended on OAL (Fig. S3C), the variation in sizes within the different stages does not allow for staging according to size.

To account for potential methodological differences (cohort analysis, kreisel tank maintenance), we compared our *in vitro* aggregate growth rates (GR_kreisel_, % d^-1^) with cohort growth rate estimates (GR_cohort_, % d^-1^) compiled from four different studies by Lüskow *et al*. (2020), all standardized to 5 °C and assuming Q_10_ = 2. Because the Arrhenius equation is not capable of generating growth rates that include negative values, we excluded chains showing potential negative growth (*n=*12) before fitting the model. The results show that estimating relative growth rates using cohort analyses systematically yields higher growth rates than those derived from measurements of individual chains (n=8) in kreisel experiments in this study and by Lüskow *et al*. (2020) (Figure 3). Whether these systematic differences are due to methodological differences cannot be said with certainty as kreisel-derived estimates are limited to only two studies and the small number of replicates (*n=*1) in our study at the first sampling time point might have led to inaccuracies in calculated growth rates. However, our kreisel-derived growth rates allowed for direct length comparisons within chains (clones), which may be the more accurate method for estimating growth rates compared to cohort analyses that take into account the average length of cohorts consisting of many chains. In summary, our results show that traditional methods of estimating growth rates may have its limitations and future research is needed to test this in more detail. Furthermore, the occurrence of negative growth rates in our data suggests that the Arrhenius equation may not be an appropriate function to account for the effects of temperature on growth rates of salps.

**Figure 3.**
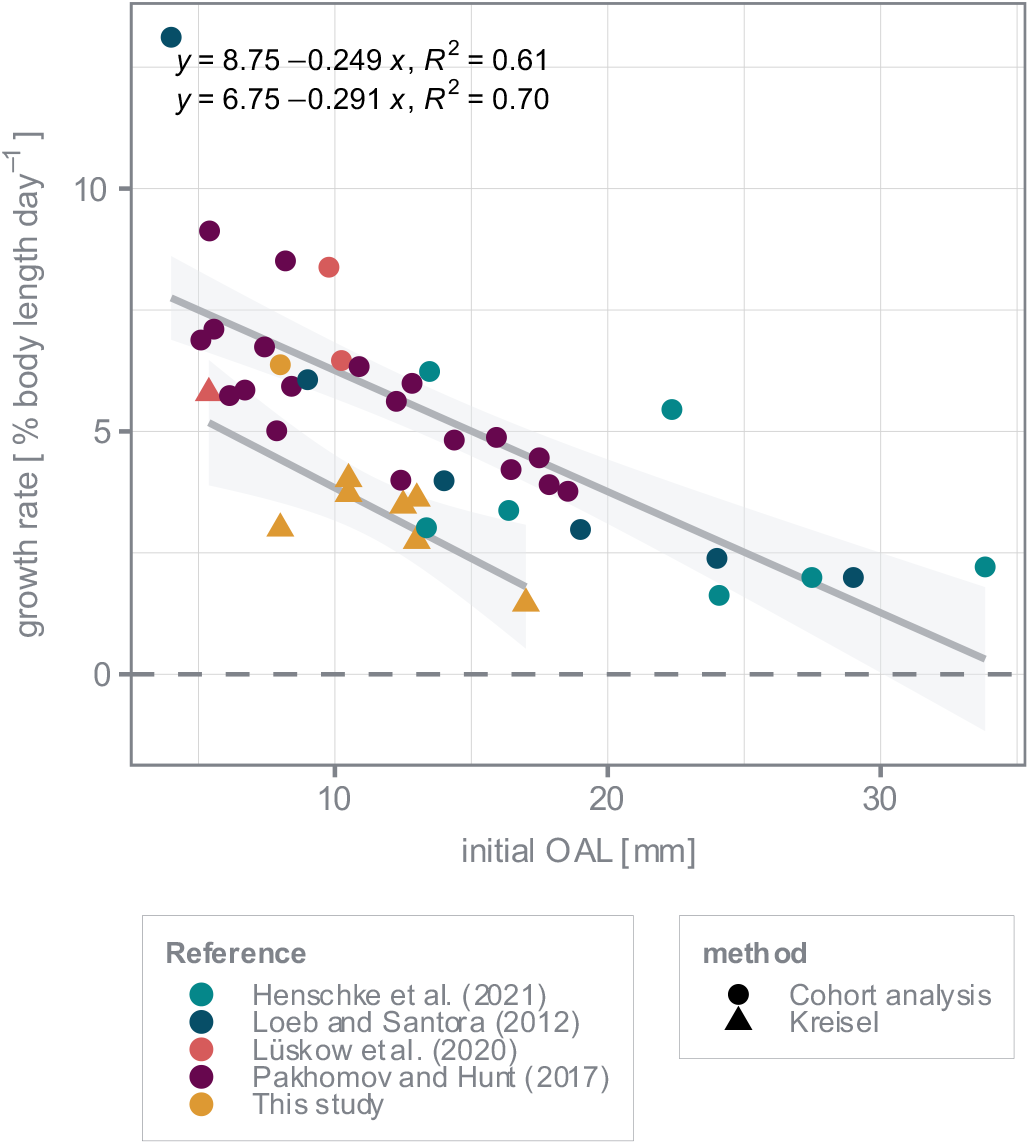
Mean growth rates of aggregates of *S. thompsoni* as a function of initial OAL [mm]. Growth rate estimates were compiled from four different studies (Loeb and Santora, 2012; Pakhomov and Hunt, 2017; Lüskow et al., 2020; Henschke et al., 2021) by Lüskow et al. (2020). Mean growth rates (for each chain) determined in this study were also added (chains n=7). All growth rates were standardized to 5 °C assuming Q10=2.

The developmental rate of aggregates was estimated by calculating the time (in days) required to develop to the next subsequent stage within each chain. Chains containing individuals of stage “X” (*n=*2, 10.53 %) were neglected in the calculation. Aggregates performed between 0 and 0.4 stage transitions d^-1^. For individuals that continued development (development rate > 0, 88.24 %) it may therefore take between 2.5 and 10 days to develop to the next development stage (Fig. S4B). It remains unresolved if the developmental rate depends on the initial stage and whether the development between stages not captured in this study would differ. Assuming, however, a general mean developmental time of 5.5 days ± 2.48 (± SD) for completing one stage transition and 6 necessary transitions from a newly released aggregate to a functional male (i.e., from stage 0 to stage M) it may take 33.00 days ± 14.86 (± SD) (range of 15-60 days) to complete the entire developmental process. This is consistent with the average time of 23 days calculated for the development of *S. thompsoni* aggregates (Lüskow *et al*. 2020). We were not able to run a similar experiment for solitaries but if we assume the same duration for the complete development of the asexual reproducing solitaries, which is defined by the development of the stolon (=chains), the complete life cycle (including sexual and asexual reproduction) would be 28-110 days, consistent with the 2-3 months proposed by Pakhomov and Hunt (2017) and much shorter than the 9 months assumed by Loeb and Santora (2012). A shorter complete life cycle (from egg to egg) of *S. thompsoni* was also suggested in individual and population modelling studies (Henschke *et al*., 2018). Our results highlight the complexity of potential life strategies of salps. However, further experiments are needed to investigate their “energetic flexibility” during different life stages.

#### S. fusiformis

Individuals of *S. fusiformis* were maintained in kreisel tanks under constant feeding and temperature conditions at the Oceanological Observatory of Villefranche-sur-Mer (France) for nine days. While aggregates from field sampling acclimated in six small kreisels for two days, solitaries were released from one chain during the acclimation period (2 days) in a separate big kreisel. Survival rates for both forms ranged from 69.1 to 85.7 % (Table 1). Our setup allowed us to track growth and development of aggregates originating from single chains (i.e., clones; asexual reproduction) and of solitaries originating from one chain (i.e., siblings).

During the nine-day incubation period, OAL of sampled aggregates was measured three times, at three, seven, and nine days after field sampling (Figure S5A). However, due to a low number of replicates on the ninth day, mean relative growth rates obtained by cross-combining individuals for each chain (*n=* 6) were calculated based on the first two samplings (sampled individuals *n=*6-7 sampling^-1^). OAL varied within chains during both samplings (± SD range 0.38-1.95 mm, Figure S5B). The mean relative growth rates for individuals cross-combined within each chain of aggregates ranged from -0.72 to 5.13 % d^-1^ (-0.10-0.68mm d^-1^) (Fig. 2B). Published growth rates of *S. fusiformis* are rare. A study by Braconnot *et al*. (1988) estimated a rate of 9-35 % d^-1^ for aggregates of *S. fusiformis* maintained in a large container (V=700 L) at 15 °C. It has to be considered, however, that these results were based on various different experiments and included very small aggregates (> 2 mm). As previously observed for *S. thompsoni,* the growth rate of *S. fusiformis* decreased moderately with increasing initial OAL (y=19-1.25x, R^2^= 0.65, Figure 2B). Therefore, the high growth rates (∼35 % d^-1^) found by Braconnot *et al*. (1988) could be related to the very small initial sizes of aggregates maintained. Two cohorts were detected in the bay of Villefranche-sur-Mer (2021). Their linear growth was 0.95 and 0.54 mm day^-1^ and their relative growth rates were 10.20 and 10.94 % day^-1^, based on initial OAL of 5.8 and 3.3 mm of aggregates, respectively (Figure S7). This growth rate is consistent with the upper end of the growth estimates for smaller individuals (∼ 10 mm) maintained in our kreisel system, confirming its favorable rearing conditions. In contrast to *S. thompsoni,* the growth rates obtained for *S. fusiformis* seemed not to depend on the initial development stage (Figure S6A). However, this can be due to the fact, that most individuals (52.17 %) were not fertilized (stage 0), limiting the number of fertilized and more advanced stages in our study. Interestingly, individuals of stage 0 showed shrinkage potential from OAL > 14 mm (Figure 2B), which may indicate, that the decrease in relative growth rates may be independent of fertilization status and is rather an obligate and/or temporal driven strategy.

On the third day after field sampling, most individuals were not fertilized yet (stage 0, 54.76 %) or just fertilized (stage 1/1.5, 45.24 %) and continued to develop during experimental maintenance (Figure S5C). Since most aggregates were still not fertilized (65%) after seven days and only some showed advanced stages (stage 3/4/4.5, 35%), we can conclude that no more salps were fertilized in our kreisel tank system. By taking the mean stage of all individuals within a chain (including stage 0) for both samplings into account for estimating the development rate, aggregates of *S. fusiformis* performed between -0.07 and 0.52 stage transitions d^-1^. For individuals that continued their development (development rate > 0, 66.67 %), it may therefore take between 1.93 and 18.67 days to develop to a more advanced stage (Figure S6B). If we assume a general mean developmental rate of 8.18 days ± 7.60 (± SD) for completing one stage transition and 6 necessary transitions from a newly released aggregate to a functional male (i.e., from stage 0 to stage M) it may take 49.10 ± 45.61 (± SD) days (range of 12-112 days) to complete the entire developmental process.

However, the large proportion of unfertilized aggregates in both sampling events in our study likely resulted in an underestimation of the calculated average development rate. Under the assumption that no more salps were fertilized in our system and neglecting individuals of stage 0 at both sampling timepoints, aggregates performed between 0.43 and 0.75 stage transitions d^-1^ resulting in a development rate between 1.34 and 2.35 days to develop to a more advanced development stage (Figure S6B). By assuming a mean developmental rate of 1.82 days ± 0.44 (± SD) for performing one transition, it may take 10.93 ± 2.69 (± SD) (range of 8-14.12 days) for full development through 6 stage transitions. This is consistent with the 14 days proposed by Braconnot *et al*. (1988) for growth and development of aggregates of *S. fusiformis* until birth of the embryo.

It is noteworthy, that most individuals (85 %) sampled in kreisel tanks on the seventh day, after field sampling showed male characteristics (development of testes). These included individuals that did not have embryos, e.g., were not yet fertilized. Aggregates are protogynous hermaphroditic, in which the ovary develops first, and the testes later (Godeaux *et al*., 1998). It has already been shown, however, that aggregates can have both, female (presence of the embryo) and male characteristics (development of testes) (Müller *et al*., 2022). It is still unclear, however, whether fertilization is a prerequisite for the development of male physiology. Our results rather indicate, that aggregates (whether fertilized or not) continued their development into males from a certain point in time onwards. This has not been described before and highlights the need to further investigate the complex reproductive cycle of salps.

Solitaries (*n=*18) were released by a chain of aggregates (*n=*21, OAL 18.17 ± 1.94 mm, stage 5) during recovery (two days) after field sampling in a big kreisel tank. Relative growth rates of solitaries maintained for 9 days at 15 °C were determined (Figure S8A). At the first sampling timepoint (day 3), solitaries were 1-2 days old and had an OAL of 14.17 ± 1.17 mm (day 3, *n=*6, Figure S8B). All solitaries sampled were at stage 1, suggesting that stolon formation began while they were still within the parent aggregate or immediately after release. The relative mean growth rates was 9.58 % d^-1^ (1.83 mm d^-1^) (Figure 2C) during maintenance of nine days. Braconnot *et al*. (1988) reported a wide growth rate spectrum of 4.8 to 40.8 % d^-1^ for solitaries maintained in aquaria at 15 °C but further comparable data are unavailable. Determining growth rates for solitary individuals represents a challenge for cohort analyses because very often solitaries are less abundant than aggregates (Lüskow *et al*., 2020). Here, solitaries accounted for a maximum of 1.72 % of total catch in field samples (mid of March 2021, Figure S7A). They were completely absent from morning samplings at the surface in the bay of Villefranche-sur-Mer during spring bloom 2021. As a result, *in situ* growth rates of solitaries could not be derived from field samplings. In our study, relative growth rates by cross-combination decreased with initial OAL, a trend that was already observed for aggregates of *S. fusiformis* (Figure S9A). After experimental day 7 (∼6 days after release by parental aggregates) almost all (83.34 %) and after nine days (∼eight days after release by parental aggregates) all solitaries (100 %) were stage 2 (Figure S8C). We can therefore assume a stage transition time between 0.16 and 0.18 stages d^-1^ and a development rate between 4.8 and 6 days (Figure S9B). If we assume 5 stage transitions (0-5), according to Foxton (1966), complete development of solitaries from release by the parent aggregate to release of the first aggregate chain itself would take 24-30 days, which is consistent with the 8-22 days proposed by Braconnot *et al*. (1988).

## Discussion

The delicate gelatinous structure of salps and their irregular seasonal and spatial occurrence make them a difficult subject for experimental research and explain the lack of any physiological data. A reliable maintenance system with specific modifications for the maintenance of salps using the well-established kreisel tank (Greve, 1968, 1975) is an advance in several respects. First and foremost, it allows the behavior of salps to be studied and to obtain reproducible physiological data such as growth and development rates. Salp growth rates are essential components in the development and parametrization of salp life cycle models (Henschke *et al*., 2018; Groeneveld *et al*., 2020). Most studies examining growth and development so far, were based on cohort analyses (Loeb and Santora, 2012; Pakhomov and Hunt, 2017; Lüskow *et al*., 2020). However, as shown in our study, cohort analyses may lead to a general overestimation of growth rates and neglect individual (and possibly stage-specific) strategies (e.g., shrinkage). Inaccuracies or high variability in input data may affect model results on population dynamics and the respective conclusions drawn from them, highlighting the need for a maintenance system-based approach. Attempts have been made in the past to maintain salps for longer periods of time, using aquaria ranging from small jars to large containers (Heron, 1972; Braconnot *et al*., 1988). However, differences in aquarium sizes, food, temperature, and sampling condition appeared to have significant impact on growth rates (Heron and Benham, 1984; Godeaux *et al*., 1998) further emphasizing the need for a standardized and thus, universal experimental system. Here, we tested and described a temperature-controlled kreisel tank system, suitable for maintaining a variety of salp species for the first time. Our results show that maintained salps have similar growth and development rates as *in situ* and therefore provide a reliable and reproducible basis for future work. Furthermore, we were able to observe key processes in the life cycle of salps, such as e.g., the timing of spermatogenesis. The hermaphroditic nature of salps, particularly the male stage, is still an under-explored area of research. A system that allows the salps to be maintained for a longer period of time is key to further study the specific characteristics of their life cycle. Additionally, manipulation of different abiotic factors (such as e.g., temperature) for various experimental designs are possible, which is important to improve existing predictions on future performance of salps, especially with respect to the possible impacts of global climate change. So far, only two studies have examined effects of increased water temperatures on salp performance using pulsation rates (Harbison and Campenot, 1979) and oxygen consumption (Trueblood, 2019) as proxies. Overall, we hope to stimulate future experimental research and thus advance the limited field of laboratory studies on salps. We also encourage others to publish detailed descriptions of their efforts and progress based on our descriptions, which is essential for comparison of the physiological data obtained here.

### Comments and recommendations

The objective of this study was to test, optimize and describe an approach for the collection and maintenance of different salp species, each characterized by different sizes, behavioral patterns, life cycle duration, and geographic distribution. We established a temperature-controlled system that meets the requirements for a successful maintenance of salps. This system is suitable for future experiments on ship-based expeditions as well as at marine research stations. Our aforementioned experiences with the system may be useful for future experimental designs, and will therefore be summarized in the following:

-*Sampling method:* Successful maintenance requires the collection of samples in healthy condition, as this directly affects the survival rate during cultivation as well as the reliability of the (physiological) parameters assessed. Towing oblique plankton nets at slow towing speed and narrow angle was already used to successfully sample pelagic tunicates in several studies (Braconnot, 1971; Perissinotto and Pakhomov, 1998; Licandro *et al*., 2006; Walters *et al*., 2019). By sampling the upper 50 m at a maximum speed of up to 2.0 knots and a closed cod end, we were able to successfully catch individuals of *S. thompsoni* in very good condition, while smaller specimens from the Mediterranean Sea rather seemed to suffer during towing (sampling depth ∼80 m). It may therefore be advisable to tow at shallow depths (max. 50 m depth) and at very low towing speeds of 1-2 knots during future sampling campaigns. In this study, the best results with Mediterranean salps were obtained by snorkeling, using beakers and zip-lock bags to catch salps directly within the water column (Figure S2). A prerequisite for this sampling technique is the close proximity of the sampling site to research facilities and the occurrence of a salp bloom, which usually occurs in spring and, to a lesser extent, in autumn. A bloom is assumed to terminate with the decline of phytoplankton concentrations, temperature change, and stratification of the water column and can last weeks to months (Menard *et al*., 1994; Licandro *et al*., 2006; Deibel and Paffenhöfer, 2009). Because of the irregular appearance of blooms and their limited time of occurrence, an onsite set up of the tank system is even more important. In our opinion, the gentlest “sampling” method is obtaining the respective form of interest via its release by the parental salp in the kreisel tanks. This reduces handling time for both generations and allows for focusing on the form of interest. This is especially important when the form is less abundant in the field as it was the case with *S. fusiformis* solitaries in the present study.
-*Number of experimental replicates:* Because even individuals of the same chain (representing clones) showed variability in growth rates in this study, a sufficient number of replicates is recommended to account for inter-individual variability and obtain statistically significant results.
-*Flow rate:* Flow should gently direct salps in a circular motion around the kreisel without causing them to tumble when passing the inflow stream. Also, flow rates should be adjusted in a way that no bubbles are formed. See also *Kreisel tank sets and modifications* for an optimal position of the spray bar and Table 1 for appropriate flow rates tested for each species and form.
-*Number of animals per tank:* The number of animals to be kept in the tanks should be carefully considered and will depend on the size and species of salps. If the tank is too crowded, salps can stick to fecal pellets, which could result in bacterial infections. Table 1 gives an indication of an appropriate number of animals per liter for the salps species investigated in this study.
-*Check animals for parasites*: Before placing the animals in the kreisel tanks, the animals should be examined for parasites (amphipods, copepods), as it has been observed that these occupy salps (Heron, 1973).
-*Establishment of non-invasive measurements through photography:* Our approach of tracking growth and developmental rate within chains and solitaries (representing siblings) allowed insights into both rates as a function of initial OAL and stage. However, length and developmental measurements were conducted after sampling individual salps. Therefore, the duration of the experiment was limited by the number of individuals available per chain. In addition, the sampling method excluded growth and developmental rate analysis of shorter chains of *T. democratica*. Therefore, future research should focus on optimizing non-invasive measurements such as e.g., photography (as in Lüskow *et al*. (2020)).

## Acknowledgments

This work was supported by project phases I+II of “The performance of krill vs. salps to withstand in a warming Southern Ocean” (PEKRIS (FKZ 03F0746A), PEKRIS II (FKZ 03F0828A)) of the German Ministry for Education and Research (BMBF). In addition, SJM received funding from the European Union’s Horizon 2020 research and innovation program under grant agreement No 730984, ASSEMBLE Plus project for travel costs, stay and access to all infrastructures at the Oceanological Observatory in Villefranche-sur-Mer (France). We thank the captain and crew of R.V. *Polarstern* (PS112) for providing salps and their excellent support during research at sea. We are also indebted to Amirhossein Karamyar and Fredy Veliz Moraleda (Alfred Wegener Institute Helmholtz Centre for Polar and Marine Research, Bremerhaven) for their advice and assistance with the kreisel tank set up. We are also grateful to the staff of the Oceanographic Observatory of Villefranche-sur-Mer for their great support during our stay in 2021 and to Alexandre Jan for the photographs of salps and field sampling.

