## Supplement_Figures for "A temperature-controlled, circular maintenance system for studying growth and development of pelagic tunicates (Salps)"

### Supplementary Material

#### Figures

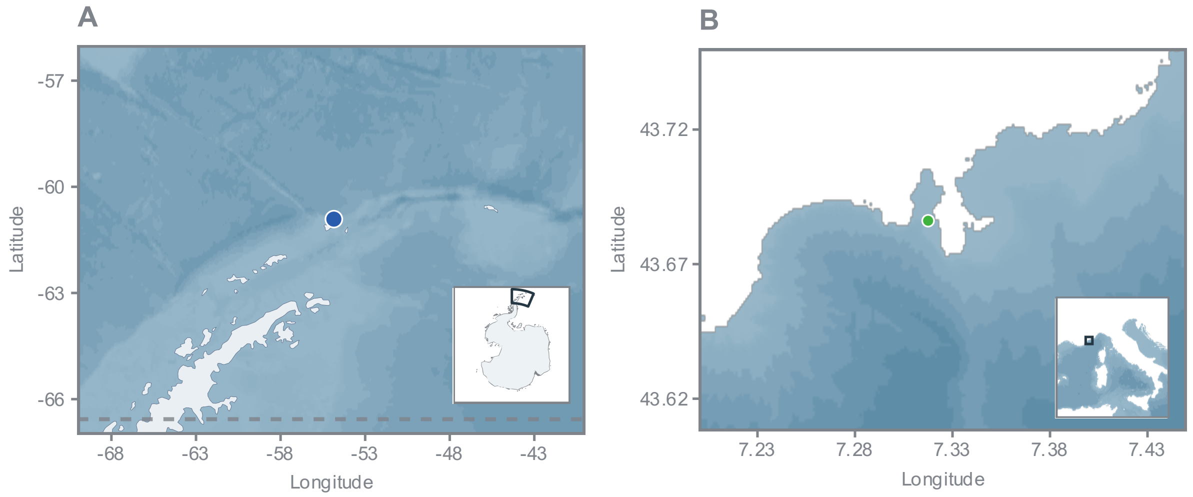


**Figure S1.** Map of sampling sites. (A) *S. thompsoni* were caught on expedition PS112 with the R.V. *Polarstern* near Elephant Island (blue circle). (B) *S. fusiformis* were sampled in the bay of Villefranche-sur-Mer, France. At the mouth of the bay, Point B, environmental data and field samples were collected to estimate *in situ* growth at regular intervals (green circle).

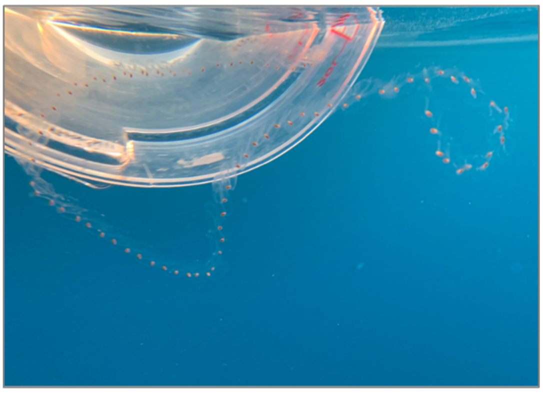

**Figure S2.** Field sampling by beaker (B). In order to catch short chains, it may be advisable to let the chain slide slowly into the beaker. For catching long chains (> 10 buds chain$^{-1}$), it is most useful to carefully pick both ends of a chain and transfer them in the bag/beacker. Photo credit: A. Jan; 2021, Oceanological Observatory of Villefranche-sur-Mer, France.

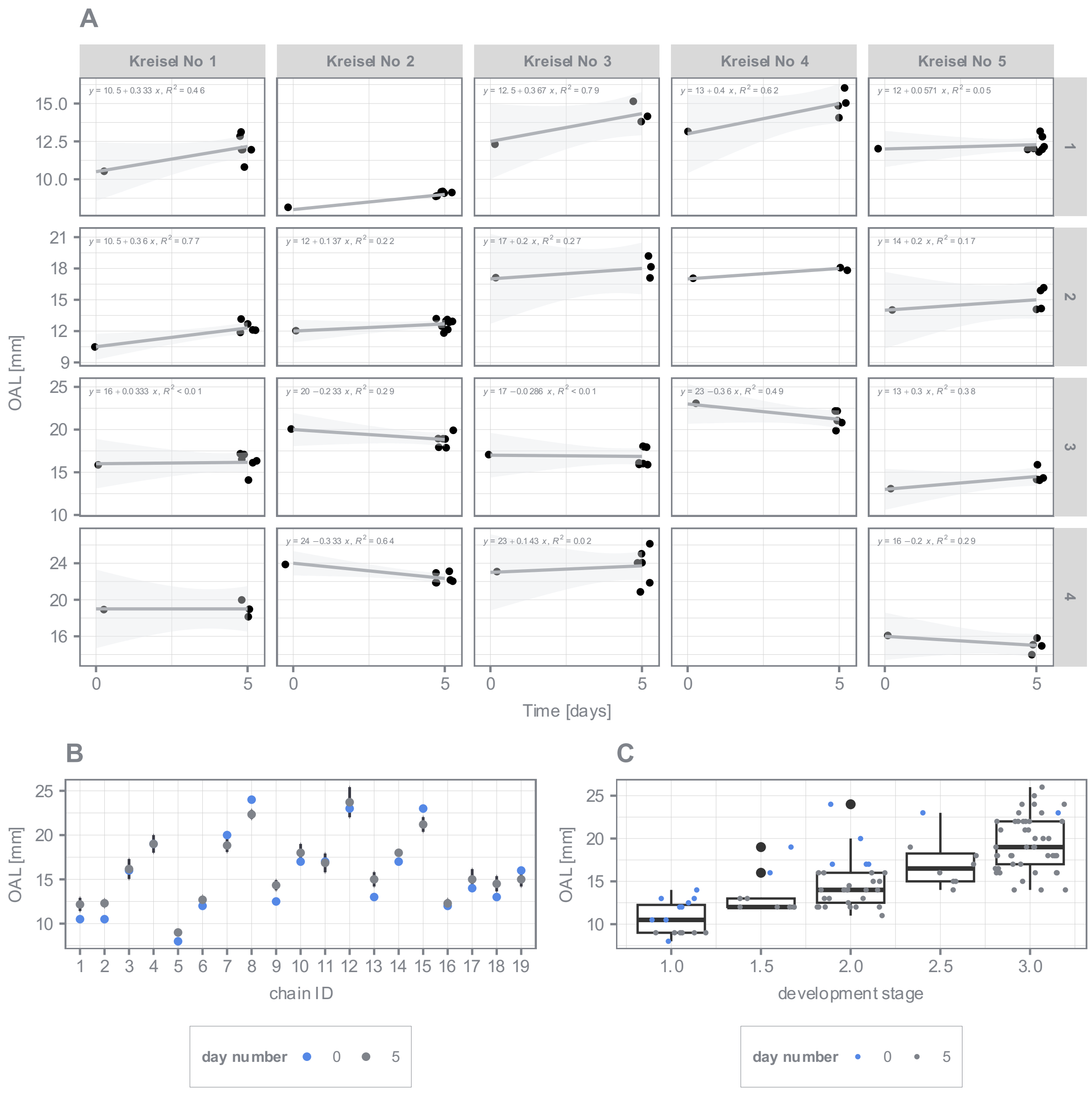


**Figure S3. Growth and developmental data of *S. thompsoni* maintained in kreisel tanks.** (A) Relationship between oral-atrial length (OAL [mm]) of sampled aggregates (n=1-8) and sampling time [days] is shown separately for each chain (total n=19) from kreisel tanks (n=5). Regression model and 95 % confidence interval were applied. (B) Mean OAL [mm] ± standard deviation for each chain and sampling timepoint (day number) are shown. (C) Boxplot of stage dependency according to Foxton (1966) on OAL [mm]. Colors indicate the numbers of days spent in the kreisel tank (0= immediately after field sampling; 5= after maintenance of 5 days in the kreisel tank).

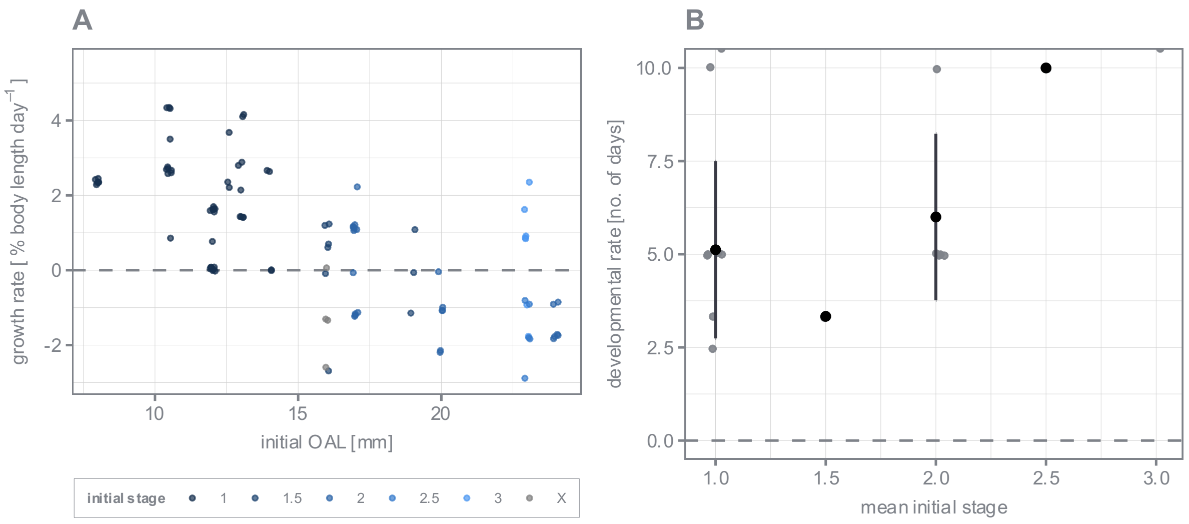


**Figure S4. Relative growth (% body length per day) and developmental rates of aggregates of *S. thompsoni*.** (A) Growth rates of individuals cross-combined were calculated as a function of initial. Different colors indicate the stages according to Foxton (1966). (B) Raw (indicated in grey) and mean and standard deviation (indicated in black) of development rates (i.e., the time it takes to transit from one stage to the next. The dashed horizontal line marks the putative threshold for the increase/decrease of daily rates.

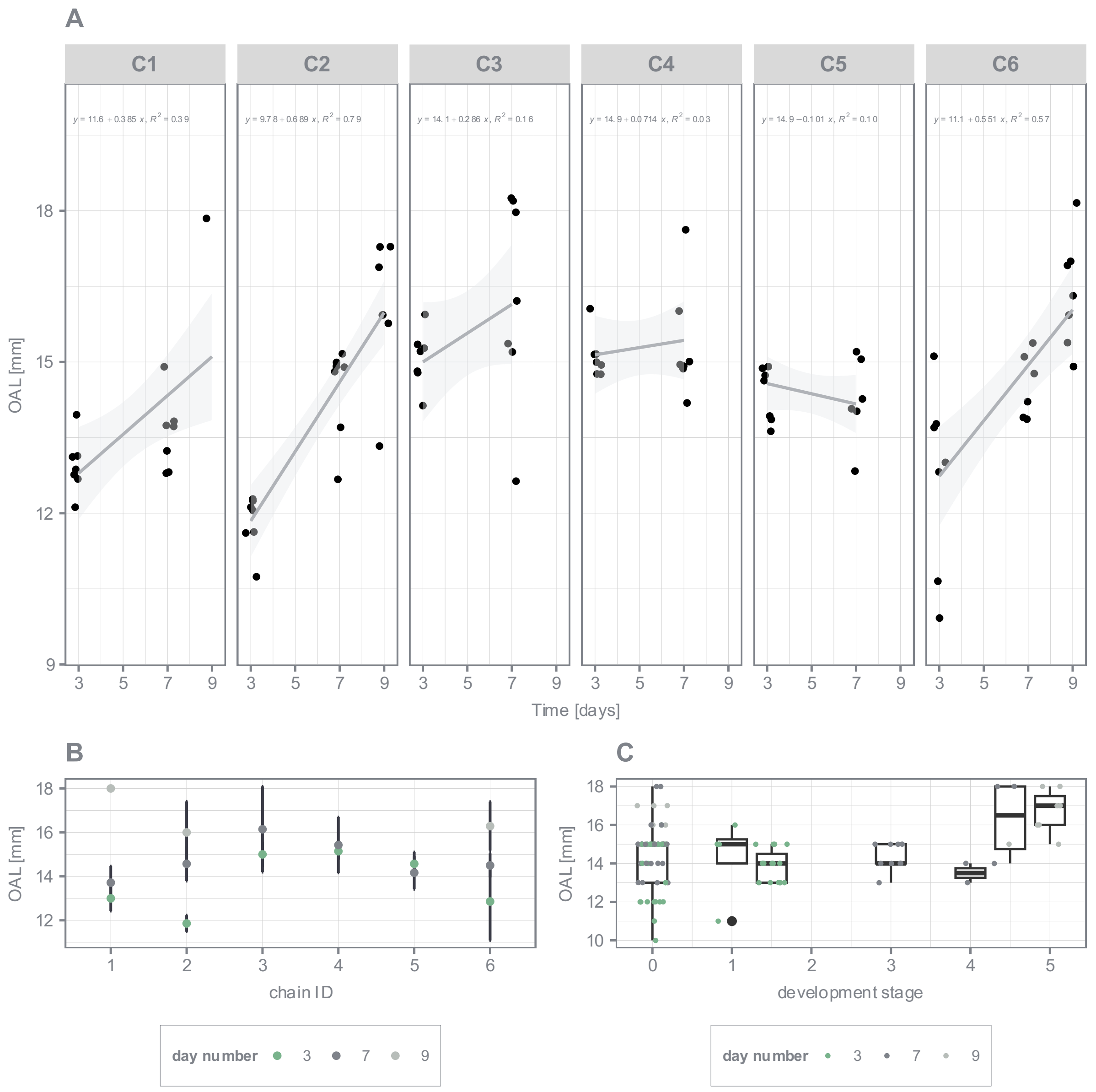


**Figure S5. Growth and developmental data of aggregates of *S. fusiformis* maintained in kreisel tanks.** (A) Relationship between oral-atrial length (OAL [mm]) of sampled aggregates (n=6-7) and sampling time [days] is shown separately for each chain (n=6). Regression model and 95 % confidence interval were applied. (B) Mean OAL [mm] ± standard deviation for each chain (n=6) and sampling timepoint (day number) are shown. (C) Boxplot of stage dependency according to Foxton (1966) on OAL [mm]. Colors indicate numbers of days spent in the kreisel tank.

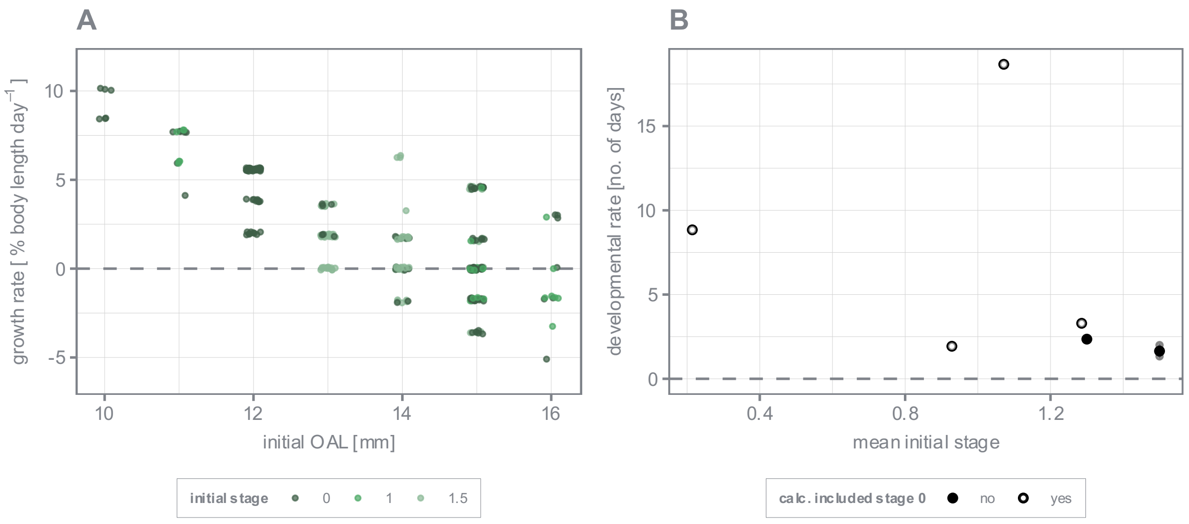


**Figure S6. Relative growth (% body length per day) and developmental rates of aggregates of *S. fusiformis*.** (A) Growth rates of individuals cross-combined were calculated as a function of initial OAL. Different colors indicate the stages according to Foxton (1966). (B) Raw (indicated in grey) and mean and standard deviation (indicated in black) of development rates (i.e., the time it takes to transit from one stage to the next) in relation to mean initial stage (according to Foxton (1966)) are shown. For aggregates of *S. fusiformis*, the developmental rate was calculated considering all stages (solid circle) and neglecting stage 0 (filled circle). The dashed horizontal line marks the putative threshold for the increase/decrease of daily rates.

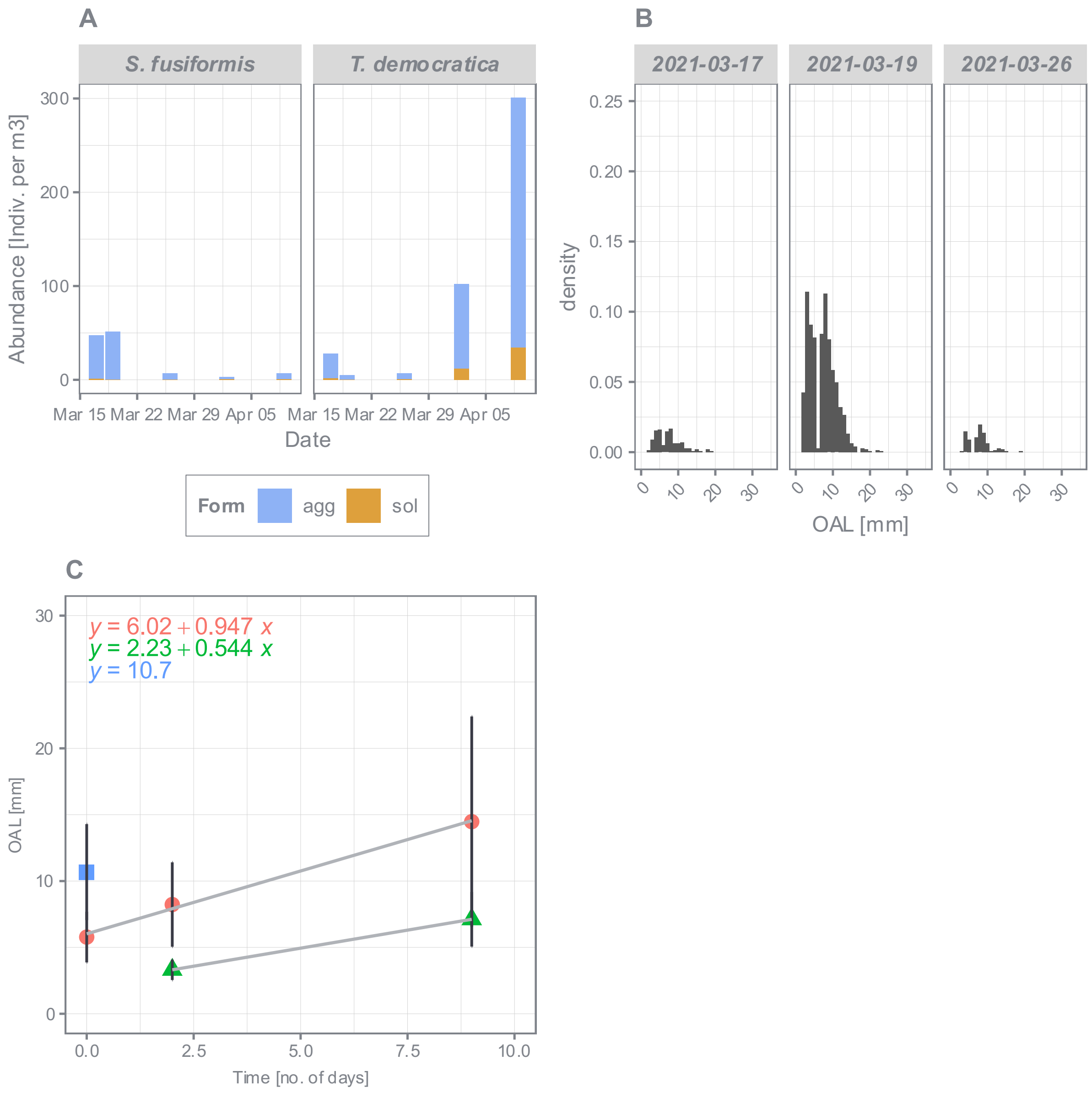


**Figure S7. Abundance [indiv/m$^3$] of salps during salp bloom in Villefranche-sur-Mer (2021) and density histogram of oral-atrial length (OAL) [mm] of aggregates of *S. fusiformis*.** (A) Abundance is given separately for *S. fusiformis* and *T. democratica* and both forms, aggregates and solitaries. (B) Density histogram of OAL of aggregates of *S. fusiformis* based on three samplings (17.03.21, 19.03.21 26.03.21). (C) Extracted mean OAL [mm] and standard deviation from cohort analysis via mixtools (Benaglia et al., 2009) were used as insert for the calculation of relative *in situ* growth rate.

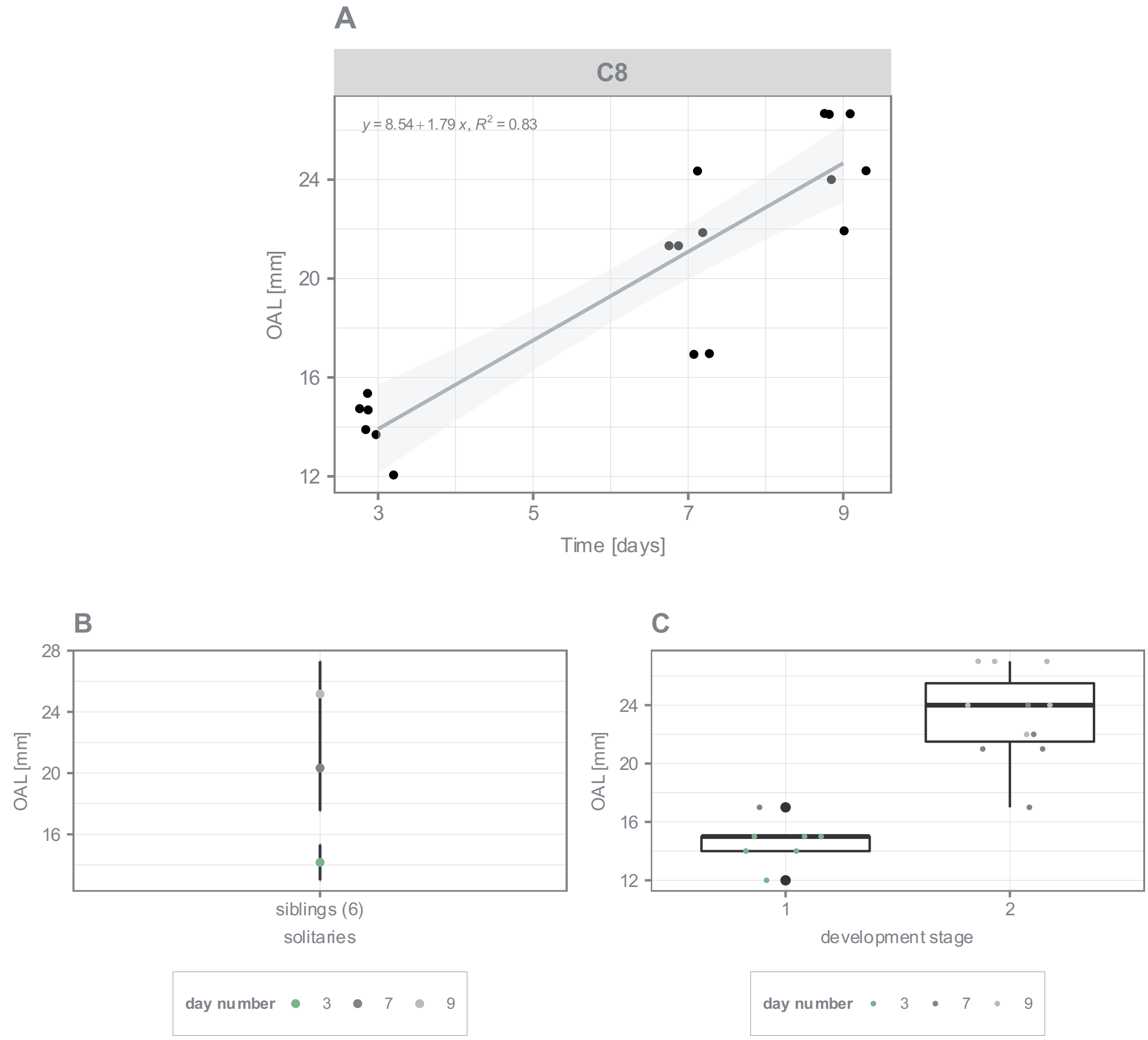


**Figure S8. Growth and developmental data of solitaries of *S. fusiformis* maintained in kreisel tank.** (A) Relationship between oral-atrial length (OAL [mm]) of sampled solitaries (n=6) and sampling time [days] is shown. Regression model and 95 % confidence interval was applied. (B) Mean OAL [mm] ± standard deviation for each group of sampled solitaries (n=6) and sampling timepoint (day number) are shown. (C) Boxplot of stage dependency according to Foxton (1966) on OAL [mm]. Colors indicate numbers of days spent in the kreisel tank.

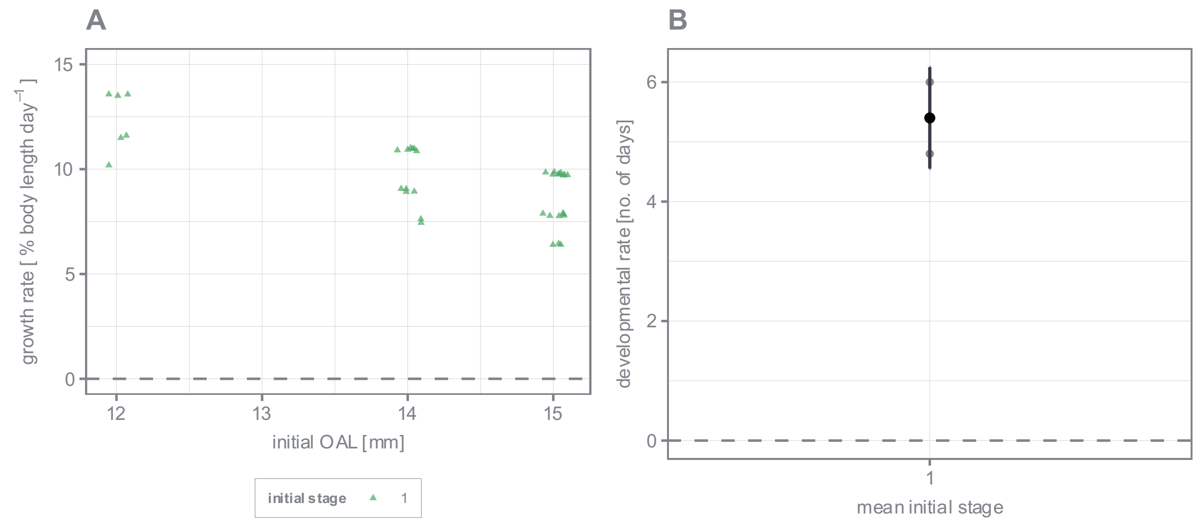


**Figure S9. Relative growth (% body length per day) and developmental rates of solitaries of *S. fusiformis*.** (A) Growth rates of individuals cross-combined were calculated as a function of initial OAL. Different colors indicate the stages according to Foxton (1966). (B) Raw (indicated in grey) and mean and standard deviation (indicated in black) of development rates (i.e., the time it takes to transit from one stage to the next) in relation to mean initial stage (according to Foxton (1966)) are shown. The dashed horizontal line marks the putative threshold for the increase/decrease of daily rates.
